# Detection and quantification of GPCR mRNA: An assessment and implications of data from high-content methods

**DOI:** 10.1101/734863

**Authors:** Krishna Sriram, Shu Z. Wiley, Kevin Moyung, Matthew W. Gorr, Cristina Salmerón, Jordin Marucut, Randall P. French, Andrew M. Lowy, Paul A. Insel

## Abstract

G protein-coupled receptors (GPCRs) are the largest family of membrane receptors and targets for approved drugs. Analysis of GPCR expression is thus important for drug discovery and typically involves mRNA-based methods. We compared transcriptomic cDNA [Affymetrix] microarrays, RNA-seq and qPCR-based TaqMan arrays for their ability to detect and quantify expression of endoGPCRs (non-chemosensory GPCRs with endogenous agonists). In human pancreatic cancer-associated fibroblasts, RNA-seq and TaqMan arrays yielded closely correlated values for GPCR number (~100) and expression levels, as validated by independent qPCR. By contrast, the microarrays failed to identify ~30 such GPCRs and generated data poorly correlated with results from those methods. RNA-seq and TaqMan arrays also yielded comparable results for GPCRs in human cardiac fibroblasts, pancreatic stellate cells, cancer cell lines and pulmonary arterial smooth muscle cells. The magnitude of mRNA expression for several Gq/11-coupled GPCRs predicted cytosolic calcium increase and cell migration by cognate agonists. RNA-seq also revealed splice variants for endoGPCRs. Thus, RNA-seq and qPCR-based arrays are better suited than microarrays for assessing GPCR expression and can yield results predictive of functional responses--findings that have implications for GPCR biology and drug discovery.

**Abstract Graphic:** 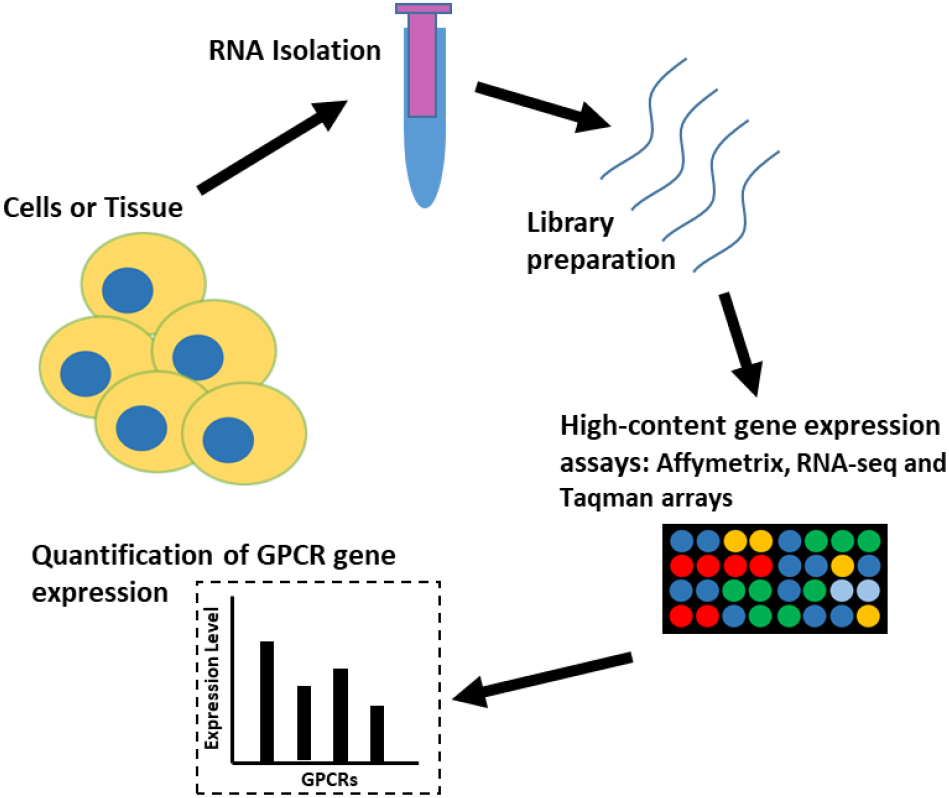

---

GPCRs, a family of >800 membrane proteins in humans, respond to a wide range of peptides, proteins, lipids, metabolites etc. and regulate a broad range of cellular processes including proliferation, metabolism, and protein synthesis. There are ~360 GPCRs that are activated by endogenous agonists, i.e., endo-GPCRs other than visual, taste and olfactory receptors. EndoGPCRs are targets for a large fraction (~35%) of approved drugs.^1^ The detection of GPCRs in cells and tissues is thus valuable for identifying GPCRs and defining their roles in cell physiology and pathophysiology as well as for identifying opportunities for drug discovery.

Detection of GPCRs by protein-based methods is challenging. Due to their low expression, GPCRs are difficult to assay by current proteomic methods plus the paucity of well-validated antibodies for many GPCRs makes it problematic to detect them by immunological techniques. As a consequence, detection of GPCRs, especially in efforts to profile their expression in cells and tissues, relies on assays of mRNA expression. Multiple methods can assess mRNA expression but their utility for defining GPCR expression has not been assessed. We thus sought to evaluate GPCR expression by parallel analysis of RNA samples from a single cell type--human pancreatic cancer associated fibroblasts (CAFs) tested with three different techniques: TaqMan arrays, RNA sequencing (RNA-seq) and transcriptomic cDNA (e.g., Affymetrix) arrays. Due to the low expression of most GPCRs, even at the mRNA level, such a comparison is important for evaluating data in public databases (e.g., CCLE^2^) that were generated using transcriptomic (Affymetrix) arrays. Because of the limited dynamic range of such arrays, it is unclear if that approach reveals accurate data regarding GPCRs. We show here that Affymetrix arrays detect fewer GPCRs than either TaqMan arrays or RNA-seq but that results from the latter two methods agree closely in terms of identity and magnitude of GPCR expression. We also provide independent qPCR validation of GPCR expression data from TaqMan arrays and RNA-seq and evidence for the predictive value of data from the latter techniques in terms of signaling and physiological response of Gq/11-coupled GPCRs.

Here, we assess GPCR expression data for the following human cells and tissues: 1) pancreatic CAFs (using Affymetrix HG U133plus2.0 arrays, Taqman arrays and RNA-seq); 2) cardiac fibroblasts (CFs), pulmonary arterial smooth muscle cells (PASMCs) and pancreatic stellate cells (PSCs) (using Taqman arrays and RNA-seq); 3) APSC-1 pancreatic cancer cell line (from CCLE, via Affymetrix HG U133plus2.0 plus RNA-seq and via Taqman arrays in our laboratory); 4) MDA-MB-231 breast cancer cell line (from CCLE via Affymetrix HG U133plus2.0 and RNA-seq); 5) Ovarian cancer (OV) tissue and Lung Squamous Cell Carcinoma (LUSC) tissue (from TCGA, via Affymetrix HGU133a and RNA-seq). This large number of sample types (also listed in **Table S1**), with data collected from different sources, facilitated a robust comparison of GPCR detection and revealed prominent differences in data generated by the different methods. These findings provide insights regarding GPCR expression by various cell types, a rationale for interpretation of mined data regarding GPCR expression, and evidence for the utility of mRNA expression data in predicting functional activity of one class of GPCRs (Gq/11-coupled GPCRs). Together such information should aid studies of GPCR biology and drug discovery.

## Methods

### Cell Culture and RNA isolation

CAFs were isolated from primary human PDAC tumors via explant and were grown as described previously^3^. At low passage (<5), CAFs were plated in 10 cm plates and grown in 5% CO_2_ at 37°C. Cells were lysed and RNA was isolated using a Qiagen RNeasy kit (Cat # 74104, Qiagen, Hilden, Germany), with on-column DNase-1 digestion (79254, Qiagen). Purified RNA had 260/280 ratios ~2 (via Nanodrop 2000c, ThermoFischer Scientific, Waltham, MA) and RNA integrity number (RIN) scores >9 (via Bioanalyzer, Agilent Technologies, Santa Clara, CA). Human fetal cardiac fibroblasts (CFs) were obtained from Cell Biologics, Cat # H6049 (Chicago, IL) and grown in 10% CO_2_ at 37°C, in low-glucose DMEM (Cat # D60646 Gibco, Dublin, Ireland) with 2% FBS (Cat # FB-02, Omega Scientific Inc, Tarzana, CA), 5 μg/L FGF2 (Cat # 130093838, Miltenyi Biotec, San Diego, CA), 5 mg/L insulin (Cat # SC360248, Santa Cruz Biotechnology, Dallas, TX), 1 mg/L hydrocortisone hemisuccinate (Cat # 07904, Stemcell Technologies Inc, Cambridge, MA) and 50 mg/L ascorbic acid (Cat # A454425G, Sigma Aldrich, St Louis, MO). Pulmonary arterial smooth muscle cells (PASMCs) were obtained from Lonza (Cat # CC-2581, Walkersville, MD) and were cultured in 5% CO_2_ at 37°C in low glucose DMEM with 5% FBS, EGF (5 ng/mL), FGF2 (5 ng/mL, Cat # 130093837, Miltenyi Biotec), insulin (same as above) and ascorbic acid (50 μg/mL). Pancreatic stellate cells (PSCs) were purchased from ScienCell Research Laboratories (Cat # 3830; ScienCell Research Laboratories, Carlsbad, CA) and cultured according to the manufacturers’ instructions.

### qPCR

RNAs were converted to cDNA via the Superscript III kit (Cat # 18080050, Thermo Fisher Scientific, Waltham, MA). cDNA was then mixed with gene-specific primers (2 μM) and Perfecta SYBR green SuperMix reagent (Cat # MP9505402K, VWR, Radnor, PA) for PCR amplification using a DNA Engine Opticon 2 system (MJ Research, St. Bruno, QC, Canada). Primers were designed using Primer3Plus software. The primer sequences are listed in Supplemental **Table S3**.

### RNA-seq

RNA sequencing was performed by DNAlink Inc (San Diego, CA for CAF samples) or the UCSD IGM core (for CF and PASMC samples), using Truseq (Illumina, San Diego, CA) stranded mRNA library preparation, with sequencing on a Nextera 500 (for CAF samples) or a Hiseq4000 (CF and PASMC samples) at 50 (CFs and PASMCs) or 75 (CAFs) base pair single reads. Data were analyzed via Kallisto v0.43.1^4^ using the Ensembl GRCh38 v79 reference transcriptome, with 100 bootstraps, to obtain transcript-level expression in transcripts per million (TPM). Kallisto bootstraps were read in R via Sleuth;^5^ GPCR data for each bootstrap were evaluated as described below to quantify uncertainty of GPCR expression quantification. Gene-level expression in TPM) was calculated using Tximpor^6^ For comparison of expression ratios of gene expression between samples, gene-level estimated counts data from tximport were input into edgeR,^7^ to obtain normalized gene expression in counts per million (CPM). RNA-seq raw data are available at NCBI GEO, at accession numbers GSE101665 and GSE125049.

### TaqMan arrays

cDNA was diluted with double-distilled H_2_O and mixed with TaqMan Universal PCR Master Mix (Cat # 4304437, Life Technologies, Waltham, MA, USA) to a final concentration of 1 mg/ml and assayed for GPCR expression using TaqMan GPCR arrays (Cat # 4367785; Life Technologies) by a 7900HT Fast Real-Time system (Thermo Fisher Scientific). Data were analyzed with RQ Manager software (Life Technologies). Gene expression was normalized to that of 18S ribosomal RNA as ΔCt; the results were consistent if normalized to GAPDH or other housekeeping genes^8^; based on previous studies, we set the TaqMan GPCR array detection threshold to a ΔCt value (relative to 18S) ≤ 25.^3,8,9^

### Affymetrix arrays

RNA samples were submitted to the Technology Center for Genomics & Bioinformatics (TCGB) at UCLA for analysis via Affymetrix HG U133plus2.0 arrays. Data as cel files were analyzed by both the MAS5 and RMA methods to quantify gene expression. Analysis was performed via the R ‘affy’ package^10^ to yield intensity estimates for each probe-set, along with detection p-values and present/absent calls (for the MAS5 method). For a GPCR to be considered ‘detected’ in Results, the Mas5 call was required to indicate ‘P’ (present) for at least one probe set for a given gene. In the event that multiple different probesets for the same GPCR indicated a ‘P’ call (with corresponding p-values < 0.05), we used the expression from the probe-set indicating highest MAS5 expression intensity, which yielded expression-ratio estimates between replicates consistent with RNA-seq and qPCR, as detailed further in Results. In general, Log2 Mas5 intensities >5 correspond to ‘present’ calls (see **Figure S3** and accompanying text on dynamic range for further details). Affymetrix data are available at NCBI GEO, at accession number GSE124945.

### Data mining

TCGA data for ovarian cancer (OV) and Lung Squamous Cell Carcinoma (LUSC) assayed by RNA-seq were downloaded from xena.ucsc.edu, as estimated gene counts and TPMs for each gene in each TCGA sample, analyzed via the TOIL pipeline.^11^ TCGA data for OV and LUSC, assayed by Affymetrix HG U133a arrays, were obtained from GEO (accession numbers GSE68661 and GSE68793 for OV and LUSC respectively) and were analyzed in R, using the methods described above for Affymetrix arrays. For computation of expression ratios between samples, estimated counts from the TOIL pipeline were input into edgeR, to obtain normalized gene expression in CPMs, allowing for comparison of expression ratios for genes between pairs of samples. RNA-seq data from the CCLE^2^ for cell lines were downloaded as gene expression in TPMs from the EBI expression atlas,^12^ from data provided on that portal, analyzed via the iRAP pipeline.^13^

### Cellular calcium assays

Intracellular calcium concentration of AsPC-1 cells was measured using the FLIPR-4 calcium assay reagent (Cat # R8142, Molecular Devices, San Jose, CA). In brief, cells were plated in black-walled clear-bottom 96-well plates overnight at ~80% confluency using media and conditions described above. Culture media was then removed and cells were incubated for 1 h at 37 °C, 5% CO_2_ in FLIPR-4 loading buffer, consisting of FLIPR-4 reagent diluted (as per the manufacturer’s instructions) in HBSS (with calcium and magnesium) buffered with 20 mM HEPES and 0.2% BSA, with pH adjusted to 7.4. Loading buffer also contained probenecid (2.50 mM, Sigma Aldrich, Cat # P8761), to prevent leakage of the calcium reagent from the cells. Calcium response was then measured via a FlexStation 3 Multi-Mode Microplate Reader (Molecular Devices). GPCR agonists were added and response in RFU (Relative Fluorescence Units) was measured over 105s for each well, yielding data for peak response and kinetics of response. For calcium assays and wound healing assays, we used the following GPCR agonists: Neurotensin (Cat # 1909, Tocris, Minneapolis, MN); 2-Thio-UTP (Cat # 3280, Tocris); Histamine (Cat # AAJ6172703, Fischer Scientific); Oxytocin (Cat # 1910, Tocris) and Sulprostone (Cat # 14765, Cayman Chemical).

### Migration/wound healing assays

Rate of migration of AsPC-1 cells was estimated using a scratch-wound assay. Cells were plated in 24-well plates and grown to approximate confluency. A scratch was made in each well using a 200 μL pipette tip, culture media was replaced to remove floating cells and the scratches were imaged using a BZ-X700 microscope. Cells were then incubated with GPCR agonists at concentrations described in the following sections and were returned to a 37 °C, 5% CO2 incubator. 24h later the same scratches were imaged once again. The area of scratches at the 0 and 24h time points was then calculated via standard protocols in ImageJ v1.52a to evaluate wound closure. Migration data were analyzed for statistical significance using Prism Graphpad (GraphPad Software, San Diego, CA) via one-way ANOVA with Tukey multiple comparison testing.

## Results

### Comparison of GPCR expression data for pancreatic CAFs

**Table S2 (top)** shows the number of GPCRs each method can detect, based on limitations of the number of primers (TaqMan arrays) or probes (Affymetrix arrays). Both methods should allow detection of a similar number of endoGPCRs. TaqMan arrays are not designed to detect chemosensory GPCRs and Affymetrix arrays also have relatively few probe-sets for chemosensory GPCRs. **Figure 1** and **Table S2 (bottom)** show that the number of endoGPCRs in pancreatic CAFs detected by RNA-seq and TaqMan arrays is greater than is detected by Affymetrix arrays. The threshold of detection used for determining whether a GPCR was detected in RNA-seq data was set to 0.2 TPM, based on analysis discussed in **Figure S1** and accompanying text. Detection thresholds for TaqMan arrays and Affymetrix arrays are discussed in **Methods**.

**Figure 1.**
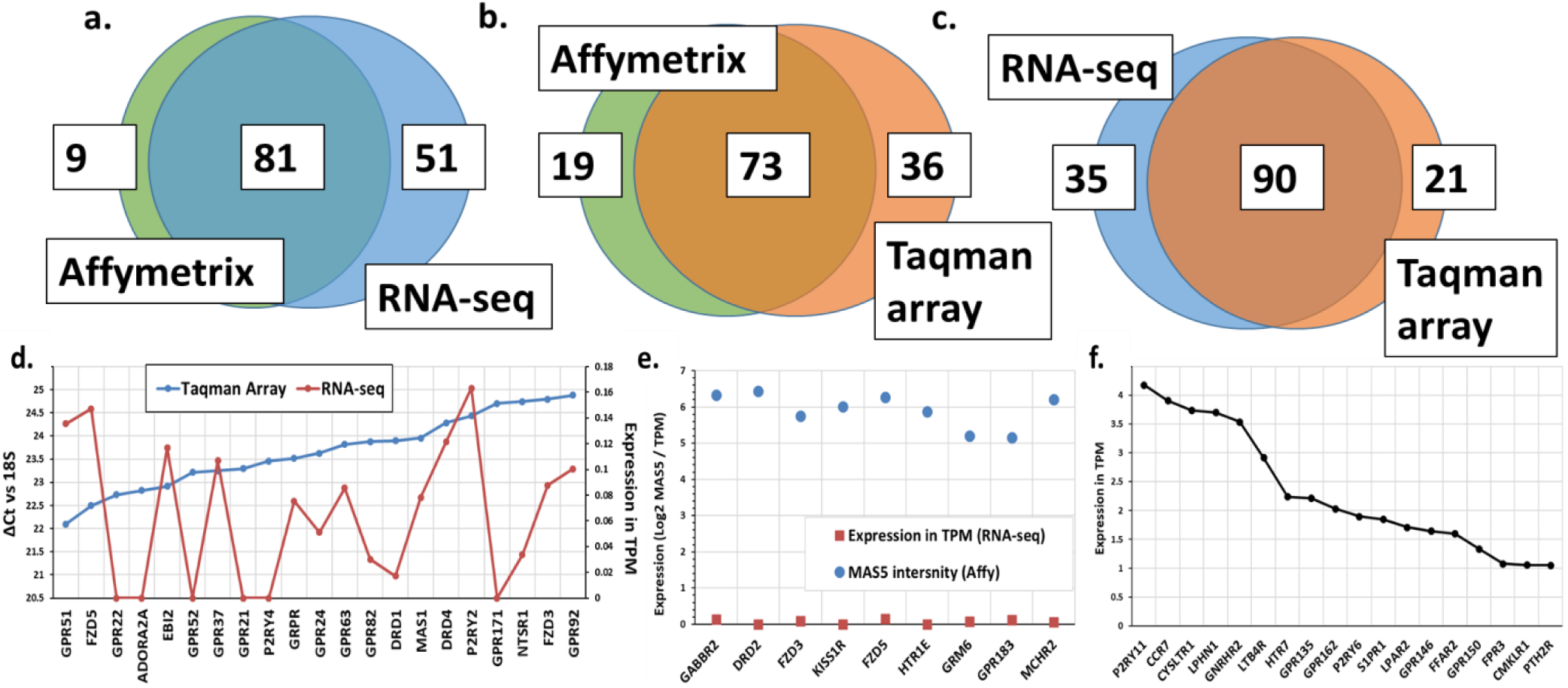
Detection of GPCRs by RNA-seq, Taqman GPCR arrays and Affymetrix (Affy) arrays. **a–c)** GPCRs for which the relevant primer probes are present by each method. Representative data are shown for an individual CAF replicate as an example; similar numbers were detected by all three methods in a second replicate. **d)** Differences in GPCR expression between RNA-seq and Taqman arrays. **e)** False positive detection of GPCRs from Affymetrix HG U133plus2.0 arrays; these GPCRs were detected by neither RNA-seq (plotted above) nor Taqman arrays (not shown) (**f**) False negatives from Affymetrix arrays that we detected by the other methods; expression is plotted for such GPCRs identified by RNA-seq

RNA-seq identified the greatest number of GPCRs, likely because this method is not limited by a fixed number of probes/primers. RNA-seq and Affymetrix HG U133plus2.0 arrays both detect a small number of chemosensory GPCRs; their level of expression was typically just above the detection thresholds defined above and in **Methods**.

Both RNA-seq and TaqMan arrays identified in common most of the detected GPCRs. Virtually all highly expressed GPCR identified by either Taqman arrays or RNA-seq were commonly detected by both, whereas Affymetrix arrays detected fewer GPCRs in common (**Figure 1 a-c**). GPCRs uniquely identified by each method were typically expressed at very low levels, i.e., near detection thresholds. Eight endoGPCRs were detected by RNA-seq but not by TaqMan arrays due to the absence of corresponding primers on the Taqman arrays.

The overall agreement between Taqman arrays and RNA-seq is further illustrated in **Figure 1 d-f**: the 21 GPCRs detected by TaqMan arrays but that were below thresholds for detection by RNA-seq were all expressed at >22 Cycles above 18S (i.e. > ~33 qPCR cycles), implying low expression levels of these receptors by either method (**Figure 1d**). Thus, all highly expressed GPCRs identified by TaqMan arrays are also detected by RNA-seq. We found a small number of apparent false positives from Affymetrix data, i.e., GPCRs not detectable by either TaqMan arrays or RNA-seq (**Figure 1e**) and numerous false negatives from Affymetrix data (GPCRs detected by RNA-seq and/or TaqMan arrays) (**Figure 1f**). The apparent false positives in the Affymetrix data were relatively low expressed (Log2 MAS5 intensity < 7). TaqMan arrays and RNA-seq show a relatively high correlation (R^2^ > 0.8) in the magnitude of GPCR expression (**Figure 2a**) but data from Affymetrix arrays correlate poorly (R^2^ < 0.4) with results from TaqMan arrays (**Figure 2b**) and RNA-seq (shown for various cells/tissues in subsequent sections).

**Figure 2.**
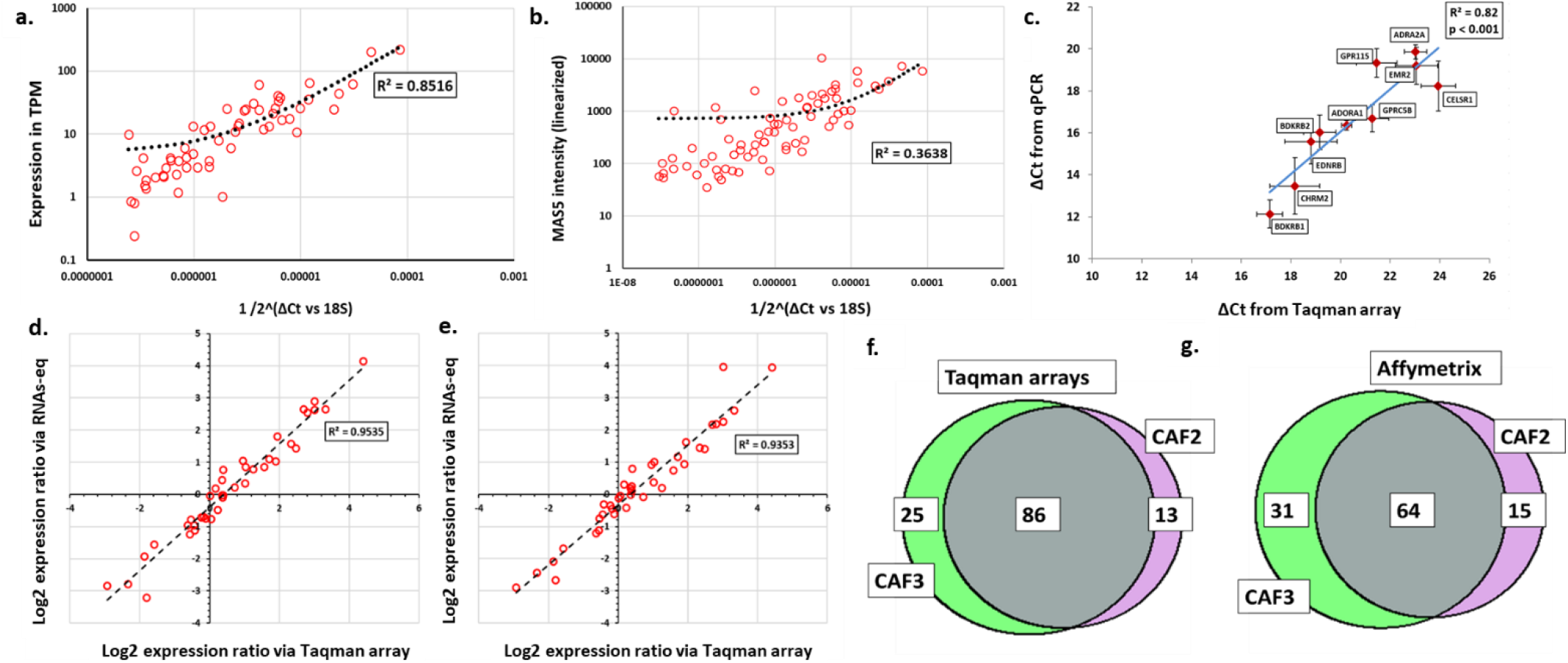
Comparison of GPCR expression levels by RNA-seq, Taqman GPCR arrays and Affymetrix (Affy) arrays with independent qPCR and comparisons of expression changes. **a)** Data from RNA-seq compared to that of TaqMan arrays; **b)** Data from Affymetrix HG U133plus2.0 arrays compared to that of TaqMan arrays. Representative data are shown for an individual CAF sample. **c)** Validation of TaqMan GPCR array data by qPCR, for N = 5 CAF samples; the data shown are mean and S.E.M. of ΔCt vs 18S rRNA. **d, e)** Correlation between expression ratios of GPCRs in two CAF samples (CAF2 and CAF3) evaluated by **d)** RNA-seq and TaqMan arrays and **e)** Affymetrix HG U133 plus2.0 and TaqMan arrays. **f, g)** Number of GPCRs in two CAF samples as detected by (**f**) TaqMan arrays and (**g**) Affymetrix HG U133 plus2.0 arrays.

Independent qPCR using SYBR green and primers designed in our lab was used to validate GPCR expression measured by the 3 methods. **Figure 2c** shows the average expression of 10 GPCRs in CAFs derived from 5 different patients, assayed via independent qPCR and the correspondence of these data with Taqman array data. We found a high degree of correspondence with results from TaqMan arrays and qPCR (R^2^ ~ 0.8) and between results from RNA-seq and qPCR (R^2^ ~ 0.8, not plotted), but not from Affymetrix arrays and qPCR (R^2^ < 0.5, not plotted).

Although Affymetrix HG U133plus2.0 arrays did not provide gene abundance estimates comparable with either RNA-seq or qPCR-based arrays, if one estimates expression ratios among biological replicates (an indication of how much a gene’s expression differs among samples), results for all 3 methods are in close agreement. **Figures 2d** and **2e** show this for ~40 GPCRs commonly detected in two CAF samples by all three methods: expression differences among samples are nearly equal for the three methods.

Are these methods equally useful for evaluating changes in expression between samples? **Figures 2f and 2g** show the overlap of detected GPCRs for two biological replicates (CAF2 and CAF3) assessed by TaqMan arrays (RNA-seq performs nearly identically^3^) and Affymetrix HG U133plus2.0 arrays. A smaller proportion of detected GPCRs was observed for n replicates tested by the Affymetrix arrays. Fewer GPCRs are consistently detectable by Affymetrix HG U133plus2.0 arrays; estimation of changes in GPCR expression is thus less feasible than with the other methods. However, for GPCRs that one can consistently quantify by Affymetrix arrays, estimates of differences in their expression are consistent with those of the other two methods. **Figure S3** (and accompanying text) shows quantitative analysis for the dynamic range of detection of GPCRs by each method, from data in CAFs. Affymetrix arrays have a narrower dynamic range than TaqMan arrays or RNA-seq, which both show very similar behavior. In addition, the correlation for all genes in general, between RNA-seq and Affymetrix arrays appears poor.

### Comparison of GPCR expression estimates in other cell types

We obtained a similarly high degree of correspondence between TaqMan array and RNA-seq data in other cell types. **Figure 3 a-c** shows the detection of GPCRs by TaqMan arrays and RNA-seq in human cardiac fibroblasts, pancreatic stellate cells (PSCs) and pulmonary arterial smooth muscle cells (PASMCs). Most GPCRs are identified by these two methods in all 3 cell types. Similar to the data from CAFs, the GPCRs commonly detected by both Taqman arrays and RNA-seq include all highly expressed GPCRs (e.g. those expressed >10 TPM). Thus, GPCR expression analysis by TaqMan arrays and RNA-seq is consistent in a variety of primary cell types.

**Figure 3.**
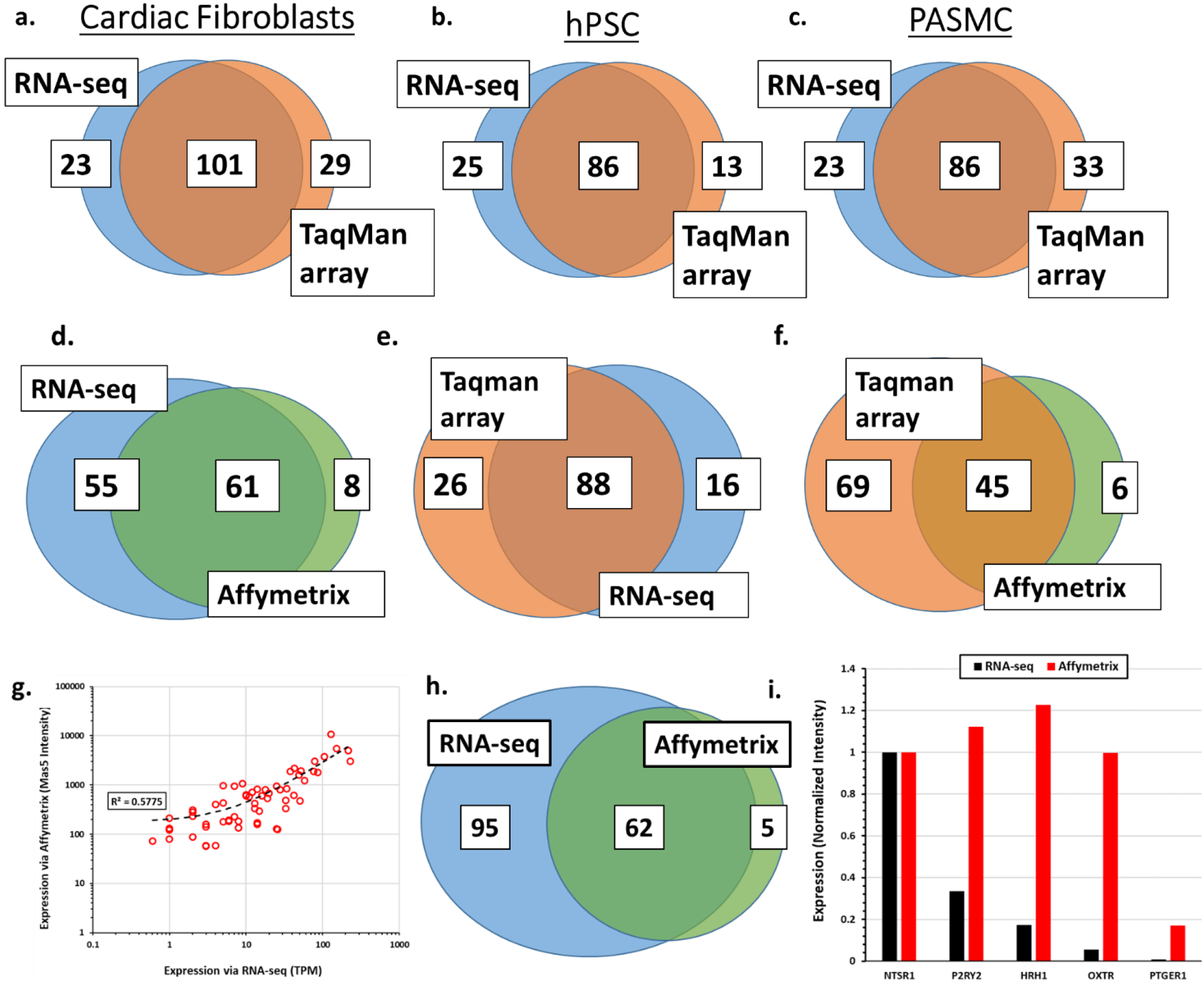
GPCR expression in other cell types. **(a-c)** Number of GPCRs detectable by RNA-seq and TaqMan arrays in individual human lines of **(a)** primary fetal Cardiac Fibroblasts, **(b)** PSCs and **(c)** PASMCs. **(d-e)** The number of GPCRs expressed by the AsPC-1 pancreatic ductal adenocarcinoma cell line (determined by Affymetrix HG U133plus2.0 arrays; CCLE), TaqMan GPCR arrays (Insel lab) and RNA-seq (CCLE and EBI). **g and h)** GPCR expression of MDA-MB-231 breast cancer cells (determined by Affymetrix HG U133plus2.0 arrays and RNA-seq; CCLE). **g)** Correlation in the GPCR detection by the two methods for the 62 commonly detected GPCRs. **h)** the number of commonly or uniquely identified GPCRs using RNA-seq or Affymetrix HG U133plus2.0 arrays. **f)** for CCLE data, expression in AsPC-1 cells of 5 Gq-coupled GPCRs tested for functional effects in **Figure 8**, linearized (for Mas5 data) and normalized to expression of NTSR1, the highest expressed of these receptors as per RNA-seq data.

To further test Affymetrix arrays with the two other methods, we assessed expression of GPCRs in AsPC-1 PDAC cells using TaqMan arrays and compared these data with RNA-seq and Affymetrix HG U133plus2.0 array data for the same GPCRs from data in CCLE^2^. We found that our TaqMan array data and the CCLE RNA-seq data showed much better correspondence than did the CCLE Affymetrix data to either of those methods/sources **(Figure 3 d-f**). Thus, GPCR expression determined by RNA-seq and TaqMan arrays assayed at different laboratories, batches of cell lines, media etc. shows greater concordance than GPCR data from RNA-seq and Affymetrix arrays, for samples prepared in the same laboratory.

We also assessed data for another cell line (MDA-MB-231 breast cancer cells) from CCLE and for tumor tissue from TCGA. We compared data from CCLE using Affymetrix HG U133plus2.0 arrays and found a similarly poor correlation between GPCR expression assayed by the Affymetrix arrays compared to RNA-seq **(Figure 3 d-e)**. Fewer GPCRs were detected by the Affymetrix arrays than by RNA-seq and the magnitudes of GPCR expression were poorly correlated (R^2^ = 0.58). **Figure 3f** shows the expression in AsPC-1 cells (from CCLE data) of 5 Gq/11-coupled GPCRs determined by RNA-seq and Affymetrix arrays; these GPCRs were subsequently studied for their functional effects **(Figure 5**). RNA-seq reveals a range of expression levels between these GPCRs, with NTSR1 and P2RY2 very high expressed while the other highlighted receptors had lower expression. By contrast, data from Affymetrix arrays implied a similar, very high level of expression for four of these GPCRs. Consistent with the RNA-seq data, data from TaqMan arrays also showed that NTSR1 and P2RY2 are especially high expressed in AsPC-1 pancreatic cancer cells (not shown), while the other receptors had lower expression.

**Figure 4.**
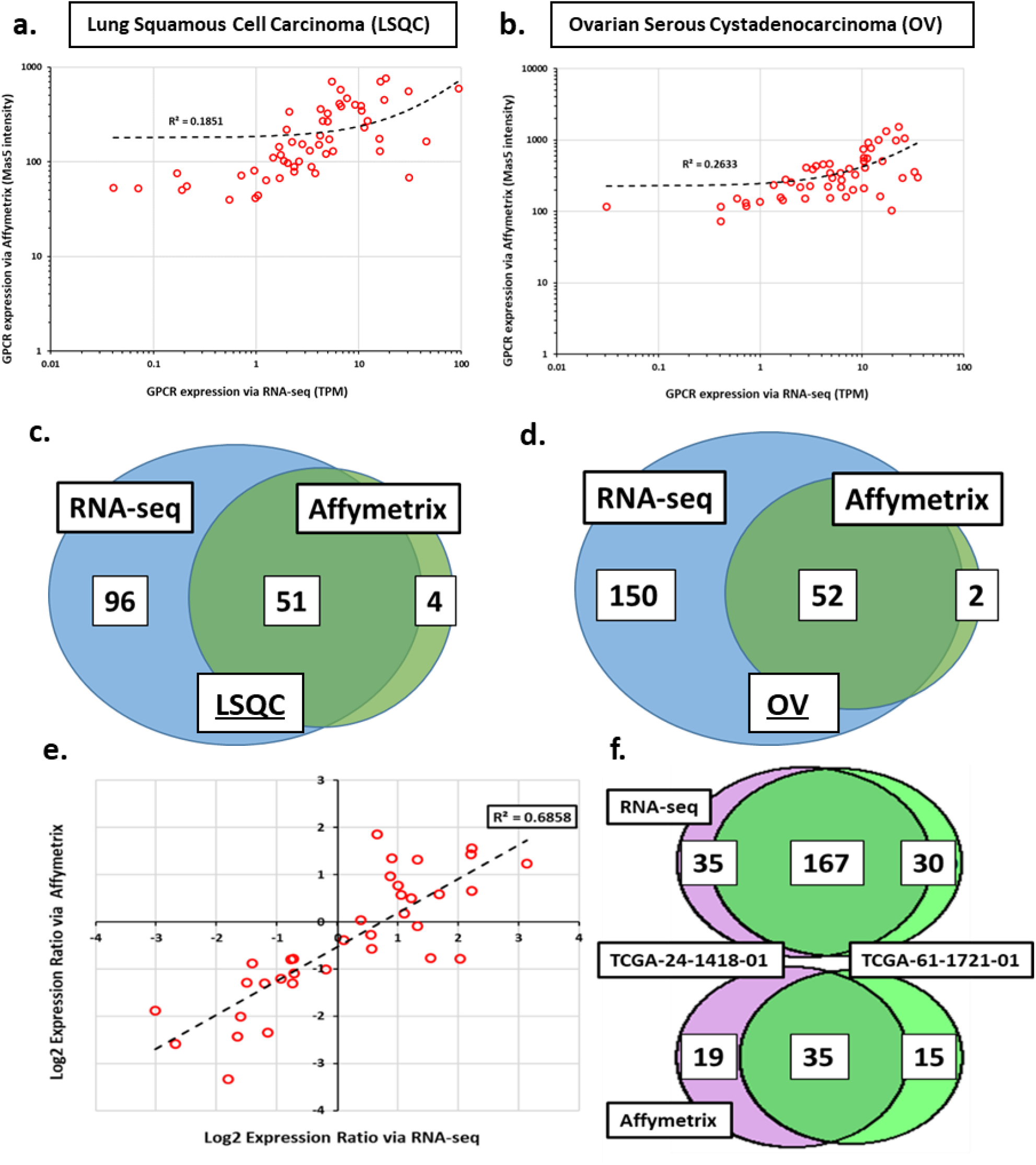
Comparison of GPCR expression data in TCGA samples generated by Affymetrix arrays and RNA-seq. **a-b)** The correlation of expression of commonly detected GPCRs and **c-d)** Venn diagrams showing the overlap in GPCR expression of randomly selected tumor samples assessed by Affymetrix HG U133a arrays or RNA-seq of ovarian cancer (OV; TCGA24-1418-01) and LUSC (TCGA-37-4141-01) tumor samples in TCGA. **e)** The correlation of expression ratios and **f)** Venn diagrams of GPCRs detected by RNA-seq or Affymetrix HG U133a array for a randomly selected pairs of TCGA OV samples.

**Figure 5.**
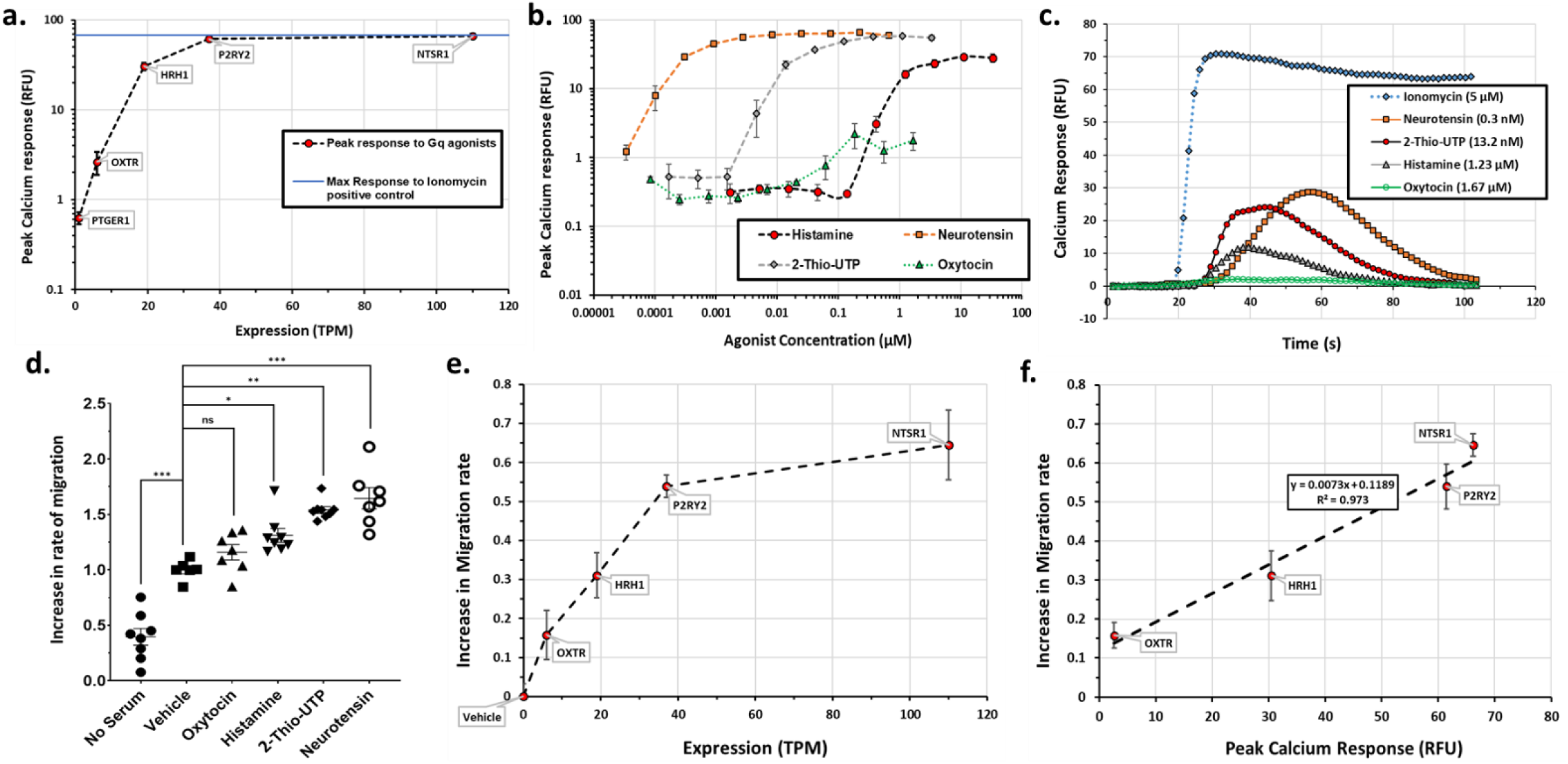
Signaling and functional response to agonists for Gq-coupled GPCRs in AsPC-1 cells. **a)** Maximal GPCR agonist-promoted increase in intracellular calcium (“calcium response”, relative to 5μM ionomycin-induced response [blue line]) for agonists of the indicated GPCRs that are expressed at different TPM in AsPC-1 cells (as determined by RNA-seq in CCLE^2^). Data shown are the mean and SEM, from 3 independent experiments. **b)** Concentration-response curves for peak calcium response by the indicated GPCR agonists compared to GPCR expression as in panel **a)**. Data shown are mean and SEM, from 3 independent experiments; **c)** Kinetics of calcium response by agonist concentrations that yield half-maximal response and kinetics of the ionomycin positive control; data shown are representative data from individual wells in a 96-well plate; other replicates showed similar behavior; **d)** Impact of treatment with GPCR agonists on the migration of AsPC-1 cells over 24 h; N ≥ 6 for each treatment. Agonist concentrations were: Oxytocin (5 μM); Histamine (10 μM); 2-Thio-UTP (0.5 μM), Neurotensin (0.1 μM); *: p < 0.05; **: p < 0.001; ***: p < 0.0001; significance was evaluated via one-way ANOVA with Tukey multiple comparison testing. **e)** The relationship between increased rate of migration and GPCR expression (as in panel **a.**); **f)** The relationship between maximal calcium response promoted by the GPCR agonist concentrations indicated in **d)** and the increase in rate of migration of AsPC-1 cells.

### Comparison of GPCR expression estimates in tumor tissue

We next tested how well RNA-seq and the Affymetrix arrays compare in assessment of GPCR expression in human tissues and in the ratios of GPCR expression in pairs of samples. For this comparison, we used gene expression data from The Cancer Genome Atlas (TCGA) for Lung squamous cell carcinoma (LUSC) and Ovarian cancer (OV) tumors. As we observed for cells, the two methods compare poorly in terms of number of GPCRs detected and magnitude of expression of individual GPCRs in the tissue samples. **Figure 4 a-d** shows the relationship between RNA-seq and Affymetrix HG U133a array data from the same tumor/donors for representative LUSC and OV samples with poor correspondence for data by the two methods (R^2^ =0.19 for LUSC and R^2^ =0.26 for OV). RNA-seq identified many more GPCRs but even for GPCRs detected by both methods, there was a poor correlation in the magnitudes of expression between the methods. The HG U133a arrays are an older, less comprehensive (i.e., smaller number of probes) product than the HG U133plus2.0 arrays we tested with CAF samples but large amounts of archived gene expression data use these or older arrays. Such archived data should thus likely be avoided for evaluating GPCR expression.

Assessment of GPCR expression between random pairs of tumor sample replicates reveals relatively poor ability to detect GPCR expression by Affymetrix HG U133a arrays compared to RNA-seq (**Figure 4 e-f**). The ratios of expression also do not correlate between the two methods, thus providing further evidence that data from older generations of Affymetrix arrays are unlikely to yield accurate data regarding GPCR expression.

### RNA-seq data suggest splice variation occurs among GPCRs

An additional advantage of the use of RNA-seq to assess GPCR expression is its ability to identify alternatively spliced transcripts, a largely unexplored aspect for GPCRs. Approximately 45-50% of GPCRs are intron-less, which may explain why splice variation in GPCRs has not been studied in detail.^14,15^ We found that many GPCRs, in particular adhesion GPCRs, which have a large number of exons, may undergo alternate splicing. **Figure S2a** shows the number of GPCRs and those that express multiple splice variant in a human pancreatic CAF sample. Nearly half of all identified GPCRs appear to show alternative splicing. As an example, ADGRE5/CD97, an adhesion GPCR, appears to have at least 5 meaningfully expressed splice variants (**Figure S2b**). These data were obtained at moderate sequencing depth; in order to more precisely identify the presence of particular splice variants in individual samples and cell types, one requires greater sequencing depth (e.g., sequencing with 150 base-pair, paired-end reads). Subsequent studies would then be needed to define biological activity of such variants. Of the three methods used here, only RNA-seq can define alternative splicing of GPCRs and with bioinformatics tools (such as Kallisto^4^) can quantify transcript levels. **Figure S2c** shows the number of detected variants for each GPCR where we noted evidence for splice variation.

### GPCR mRNA expression shows concordance with signaling and functional response

To validate the GPCR expression data, we tested whether the level of mRNA expression can predict GPCR response with respect to signaling and functional activities. We opted to study a set of GPCRs which couple to Gq/11 Gα proteins, thus signaling via increases in intracellular calcium: NTSR1 (the Neurotensin receptor), P2RY2 (the P2Y2 purinergic GPCR), HRH1 (the Histamine H1 receptor), OXTR (the Oxytocin receptor) and PTGER1 (the EP1 prostaglandin receptor), spanning a range of expression (assayed via RNA-seq in CCLE^2^) from >100 TPM (NTSR1) to 1 TPM (PTGER1) in AsPC-1 pancreatic cancer cells (**Figure 5A**). All 5 GPCRs have well-known agonists that act via Gq/11^25^: Neurotensin (NTSR1); 2-Thio-UTP (P2RY2), Histamine (HRH1), Oxytocin (OXTR) and sulprostone (PTGER1). Each agonist should activate the specified receptor due to their known pharmacology and based on the GPCRs expressed in AsPC-1 cells. For example, HRH1 is the only HRH receptor, OXTR is the only vasopressin and oxytocin receptor family member expressed and among targets for sulprostone, PTGER1 is the only one expressed in AsPC-1 cells. Similarly, P2RY2 is the only purinergic GPCR expressed, which is activated by 2-Thio-UTP.

We first tested the calcium response (i.e., the increase in cytosolic calcium) in AsPC-1 cells for each agonist, over a range of concentrations. **Figure 5a** shows the peak (“maximal”) calcium signal recorded for each ligand (at saturating concentrations [**Figure 5b**]) as a function of the magnitude of expression of its cognate GPCR target. We observed a sigmoidal behavior, wherein the calcium response plateaus for highly expressed GPCRs; the maximal signal was equal to that elicited by ionomycin, a positive control. PTGER1 did not elicit a signal, implying that 1 TPM may be a threshold for detection of Gq/11 signaling in these cells by this method. **Figure 5b** shows the concentration-response for each agonist that increases cytosolic calcium. The apparent EC_50_ values for these ligands vary somewhat from values in the literature, for example, the EC_50_ for neurotensin signaling at NTSR1 is ~0.3 nM, approximately an order of magnitude lower than that reported in the literature^25^. This raises the possibility that signaling in native cells may differ from that in model systems where such data are typically generated. The kinetics for ionomycin and for agonists at concentrations approximately corresponding to half-maximal response (**Figure 5c**) suggest differences in the rates of activation of the different GPCRs, although the calcium transient is ~ 60-90s in all cases.

We next tested the ability of these agonists (excluding sulprostone, as we obtained no evidence of calcium response for this compound), to stimulate migration (using a wound-healing assay) at concentrations that correspond to maximal activation of their respective GPCRs. **Figure 5d** shows the effect of agonist treatment on the rate of wound-closure, compared with vehicle-treated cells. We used cells plated without serum as a negative control. Higher GPCR expression and stronger calcium response results in greater stimulation of migration (**Figure 5e and f**), thus allowing us to relate GPCR expression, signaling and functional response. Based on data in **Figure 3f**, Affymetrix data in the same cell line, from the same source (CCLE) failed to resolve the differences in expression between these GPCRs that were observed via RNA-seq. As a consequence, the data from the Affymetrix arrays did not permit accurate identification and stratification of GPCRs based on their expression and hence those results could not reliably predict which GPCRs were highly enough expressed to yield a strong functional response. Thus, data from Affymetrix arrays do not reliably identify the GPCRs that are most highly expressed. In screening for expression of GPCRs for subsequent drug/target discovery studies, Affymetrix arrays should likely be avoided in favor of other methods.

## Discussion

Comparison of three methods for high content screening of GPCR mRNA expression reveals that TaqMan GPCR arrays and RNA-seq show comparable performance, whereas Affymetrix arrays perform at a lower level. A likely reason for the latter result is the generally low expression of GPCRs, such that they are outside the dynamic range for optimal detection by arrays designed to assess the entire transcriptome. Highly expressed GPCRs, especially ones that are well-characterized, are generally reliably detected by either TaqMan GPCR arrays or RNA-seq. By contrast, Affymetrix arrays fail to identify large numbers of GPCRs and show poor correlation of expression and estimates for changes in expression compared to the other two methods. GPCRs, as a large family of genes, with a large (3 orders of magnitude) range of expression provide additional support for the accuracy and completeness, especially of RNA-seq data, and complement other validation studies.^16^ The numerous false negatives observed with Affymetrix data are likely attributable to lower sensitivity, i.e., for expression thresholds < 4-5 TPM in corresponding RNA-seq data, GPCRs will frequently be undetected by Affymetrix arrays.

Comparison of the three methods (**Table 1**) shows that TaqMan arrays or RNA-seq are preferable for GPCR detection and profiling. TaqMan arrays require minimal bioinformatic effort and thus can rapidly generate data. Important advantages of RNA-seq include: 1) the number of GPCRs that are potentially detectable (**Table S2**), since gene-specific primers or probes are not needed and 2) RNA-seq detects non-GPCR genes (including data for post-GPCR signaling components), yielding far more information than do TaqMan arrays. RNA-seq thus has the potential to explore other aspects of GPCR biology, such as pathways for cellular regulation.

**Table 1.** Comparison of three high-content assays for identifying and quantifying GPCR expression.

|  | **<u>TaqMan GPCR arrays</u>** | **<u>RNA-seq</u>**<br>**(75 bp, single reads,**<br>**~25\*10<sup>6</sup> reads/sample)** | **<u>Affymetrix arrays</u>**<br>**(HG U133 plus 2.0)** |
| --- | --- | --- | --- |
| Ability to detect genes | Most endoGPCRs, but very few other GPCR types | All annotated GPCRs (~800) and other genes in the reference genome used | Most endoGPCRs, but a relatively small proportion of chemosensory GPCRs |
| Quality of gene expression quantification | High sensitivity, with repeatable, robust results | High sensitivity, with repeatable, robust results | Poor sensitivity, large numbers of false negatives |
| Assay Cost* | ~400 \$ / sample | < 200 \$ / sample | ~450 \$ / sample |
| Analysis requirements | RQ manager (Invitrogen) | Relatively complex multi-step analysis pipeline using multiple tools | Standard algorithms such as MAS5/RMA, implementable in R |
| Recommended amount of input (total) RNA | ~ 1000 ng | ~ 200 ng | ~ 1000 ng |
| Other factors | Normalization requires use of housekeeping genes, expression quantified as $\Delta C_t$ vs housekeeping gene | Normalization is independent of housekeeping genes, gene expression quantified in units such as CPM/TPM | Normalization is independent of housekeeping genes, expression quantified via MAS5/RMA intensity |
| Data Mining for GPCR expression | Not widely available, but where such data are provided, should be usable | Usable, with appropriate attention to data normalization and standardization | Mining of archived data, especially for older Affymetrix arrays not advised for GPCR expression |
| **<u>Overall assessment</u>** | Recommended for focused study of GPCRs, especially for time-sensitive data; easy data analysis | Recommended for study of GPCRs but requires access to bioinformatics tools, plus may be time-lag in generation of gene expression data | Not recommended to assess GPCR expression |

\* Costs shown are with academic pricing, these are subject to variation, depending on access to core facilities as well as institution type.
**Abbreviations:** CPM (Counts per Million), TPM (Transcripts per Million), ΔCt (Delta/difference in Cycle threshold).

As a consequence of the ongoing decrease in cost of sequencing, the potentially lower expense to conduct RNA-seq is another advantage of this technique. A “hidden” cost of RNA-seq involves data analysis and storage, which can substantially increase the expenditure for RNA-seq. RNA-seq requires time for library preparation and sequencing, as well as bioinformatic analysis. By contrast, data analysis of qPCR-based arrays can be done quickly. Other qPCR-based arrays are available that may yield comparable data but at a lower cost than TaqMan® arrays, e.g., SYBR green-based arrays (e.g. Cat # 10034500 [Bio-Rad] and Cat # PAHS-071Z [Qiagen]).

A further advantage of RNA-seq is that it has become a method of choice for many large databases and consortia (e.g., TCGA and GTEx^17^). The abundance of such publicly available RNA-seq data facilitates data mining and yields information for new studies, including with respect to GPCR signaling components and how GPCRs and such components may have altered expression profiles during physiologic perturbations and in disease states. Limited public data are available for GPCR arrays so mining of such data is less feasible.

Affymetrix array-derived data is found in sources such as Gene Expression Omnibus (GEO) and other databases and for many years has been used for data mining. The findings here suggest that for GPCRs, mining of Affymetrix array data is not advisable. Moreover, to the extent that GPCRs may be differentially expressed in cells or tissues, such as in disease,^3^ the large number of false negative results from Affymetrix arrays may impact on other analyses, such as in pathways and networks, in which GPCRs may be involved. The inferior dynamic range of Affymetrix arrays is likely not limited to GPCRs and may impact on the detection of other low expressed, but functionally important, genes. Thus, caution is advised in the use of Affymetrix arrays and mining of Affymetrix array data.

The study of the ‘GPCRome’ and individual GPCRs identified by the methods compared here has the potential to yield important insights regarding the regulation of cells and tissues in health and disease. Moreover, GPCRs are targeted by ~35% of FDA- and EMA-approved drugs^1^ and represent the largest family of drug targets. Given their high druggability, identification of GPCRs in novel contexts can aid in drug discovery efforts.^18^ The current results and other data show that cells express large numbers of (>100) GPCRs; these include GPCRs targeted by approved drugs, orphan receptors and many GPCRs for which tool compounds (but not approved drugs) may exist. Multiple studies of ‘GPCRomics’ combined with signaling and functional analyses have revealed novel roles for GPCRs in numerous cell types.^3,8,19–22^ A growing number of studies have also begun to reveal the extent to which the presence of splice variants among GPCRs may impact their functional activity.^14,15^ Consequences of such alternative splicing include the presence of receptor isoforms with altered ligand binding (primarily due to changes in the N-terminus), altered downstream signaling (including ‘decoy’ receptors that bind ligands but have no functional activity), and potential effects on receptor trafficking, internalization and localization within specific cellular domains. This remains a largely understudied aspect of GPCR biology. The presence of numerous GPCR splice variants at the mRNA level underscores the need for further investigation.

Because mRNA expression may not necessarily predict protein expression, we undertook functional studies of Gq-coupled GPCRs to test the concordance of mRNA expression with signaling/functional data. In general, for GPCRs, direct measurement of protein expression has been challenging, due to a) difficulty in obtaining well-validated antibodies and b) the low magnitude of expression of GPCRs. Thus, indirect methods to verify protein expression are needed. Here, we show that GPCR mRNA expression predicts signaling and functional response for multiple Gq/11-coupled GPCRs in a pancreatic cancer cell line. Highly expressed GPCRs (e.g., NTSR1 and P2RY2) which are among the 10 most highly expressed GPCRs overall in these cells, show very strong agonist-induced increases in intracellular calcium response and prominent functional response (migration), both of which appear to saturate at high levels of GPCR expression. We are not aware of prior such data for Gq/G11-coupled GPCRs, in particular in native cells.

Other results that imply a concordance between mRNA expression and functional response of prostanoid receptors in fibroblasts^23^ and adrenoceptors in IPS-derived cardiomyocytes^24^ support this observation in other GPCR systems. Thus, encouraging initial data (including those shown here) suggest that GPCR expression data can provide a useful first step in drug/target discovery efforts, with highly expressed GPCRs (or differentially expressed GPCRs in disease) likely to be the favored candidates for subsequent validation. Further studies are needed to determine the extent to which GPCR expression predicts intensity of downstream signaling events, such as protein phosphorylation, transcriptional regulation, etc. Affymetrix arrays are not recommended for such efforts, as they do not adequately distinguish between high-expressed and low-expressed GPCRs and, in addition, fail to detect many GPCRs.

Omics data such as those presented in this study reveal the high expression of numerous orphan GPCRs in human disease, for which such validation studies are more challenging. Given the apparent concordance between magnitude of expression and functional response of GPCRs, such data highlight specific highly expressed orphan GPCRs as priority candidates for further study and for attempts at deorphanization.

The data presented here highlight aspects of GPCR biology that merit further study. These include: a) what is the functional impact of alternative splicing of GPCRs? b) the high expression of many orphan GPCRs underscores the importance for further deorphanization efforts; the physiological role of much of the GPCRome remains unknown; c) given the abundance of GPCRs expressed in different cell types, do GPCRs that are more widely/ubiquitously expressed than others have a functional significance? d) which mechanisms regulate the expression of individual or groups of GPCRs in particular cell types? We anticipate that GPCRomic efforts, in combination with other techniques, will help address such issues, advance understanding of GPCR biology and aid in efforts to develop novel therapeutics.

## Abbreviations

CPM: Counts per Million
TPM: Transcripts per Million
ΔCt: Delta/difference in Cycle threshold

## Acknowledgements

Studies in our laboratories related to this topic were supported by research and training grants from the National Institutes of Health (CA121938, HL007444, CA189477, AG053568) with additional research support from the Department of Defense (W81XWH-14-1-0372), Bristol Myers Squibb, an ASPET David Lehr Award and the Padres Pedal the Cause PTC2017 award.

Data generated by the TCGA (The Cancer Genome Atlas) Research Network were used in this study.

