## Supporting Figures and Text for "Detection and quantification of GPCR mRNA: An assessment and implications of data from high-content methods"

### Contents

**Supplemental Figures and Text**

- **3 figures, Figures S1-S3**

**Supplemental Tables**

- **3 tables, Tables S1-S3**

**Number of Pages: 9**

### Supplemental Figures and Text

***Setting a threshold for detection of GPCRs in RNA-seq data***

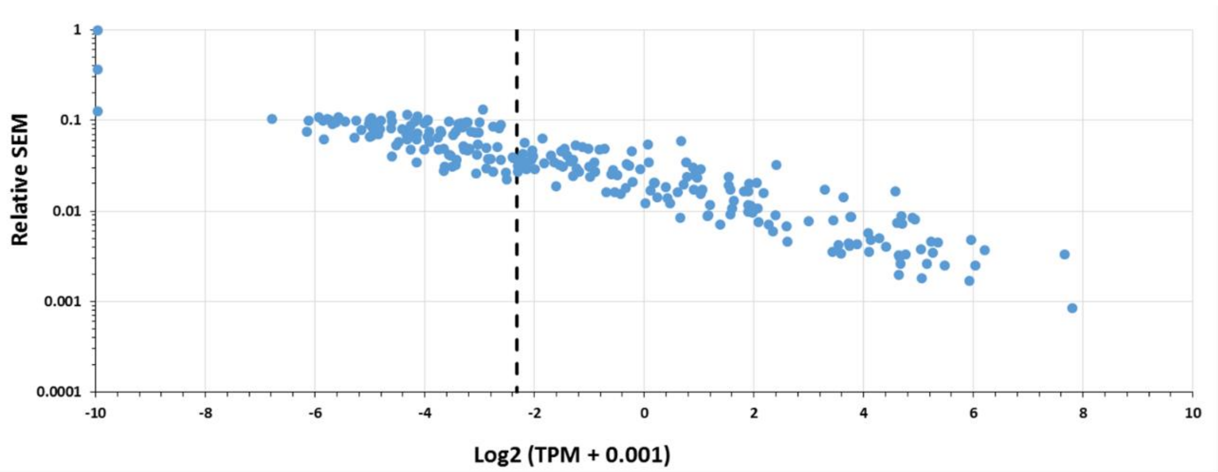


**Figure S1**. **Setting a detection threshold for RNA-seq.** In general, a threshold of >0.2 TPM restricts 'detected genes' to those with relative SEM < 10% at the indicated sequencing depth.

***Setting a threshold for detection of GPCRs in RNA-seq data***

Gene expression by RNA-seq was estimated using Kallisto, with n = 100 bootstraps to arrive at estimates for transcript-level expression (and uncertainty of quantification) in TPM, and via the expectation maximization method to arrive at an optimal estimate of transcript abundance/expression. The Kallisto bootstraps allow for quantification of the uncertainty in quantification of transcript expression, which is utilized by the Sleuth tool[5] to evaluate differential expression. Gene-level expression in TPM and gene-level counts data were obtained via Tximport[6]. In addition, gene-level estimates (in TPM) from each bootstrap were estimated by summing abundance in TPM for all transcripts mapping to each gene. These gene-level abundance data from each bootstrap allowed us to estimate the mean bootstrap-estimated expression in TPM (which is typically very similar to the 'optimal' estimate from the Expectation-Maximization method + tximport) while also allowing inference of the relative error of such estimates, based on the variance among the 100 separate bootstrap abundance estimates. **Figure S1** shows the relative error for each GPCR gene across 100 bootstraps, plotted against mean expression across all 100 bootstraps; genes expressed above ~0.2 TPM have relative errors <10%. As the magnitude of gene expression decreases, the smaller number of available reads for a given gene makes estimates of gene expression more uncertain and hence the estimation error increases. We thus chose an expression threshold of

0.2 TPM for our RNA-seq data. At higher expression levels, the bootstrapping method suggests a standard error of <10% in abundance estimates at ~25 million 75 bp single reads per sample; greater sequencing depth should lower this detection threshold. As shown in results below (**Figure S3**), the detection thresholds selected for Taqman arrays and RNA-seq result in identical dynamic ranges for detection of GPCRs.

***RNA-seq data suggest splice variation occurs among GPCRs***

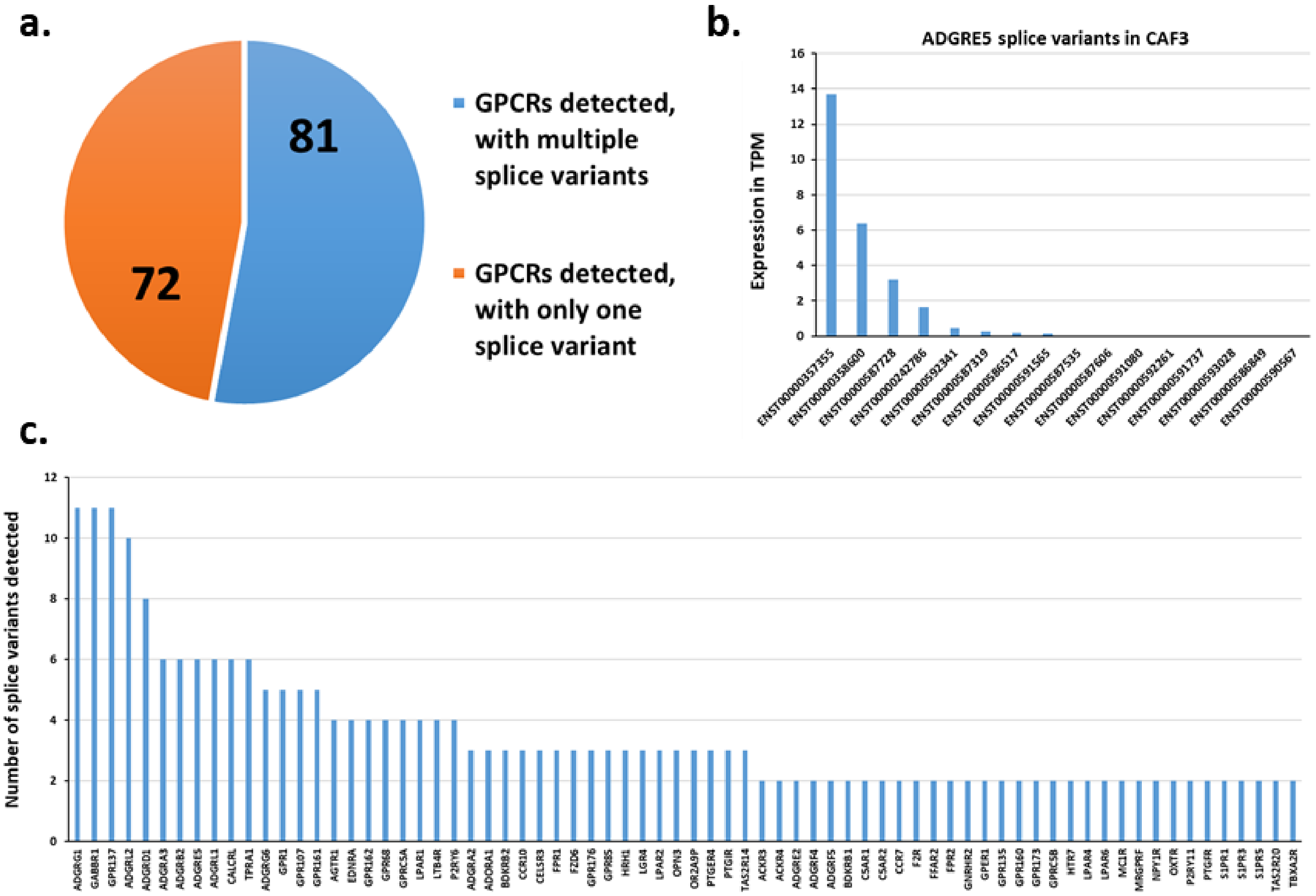


**Figure S2.** Expression of splice variants of GPCRs in CAFs: CAF3 as an example. **a)** The number of GPCRs detected with multiple transcripts expressed at > 0.2 TPM expression threshold; **b)** As an example, the expression of different transcripts for ADGRE5 (aka CD97). **c)** GPCRs with multiple transcripts detected and the number of transcripts expressed at >0.2 TPM for each GPCR.

***Comparison of the dynamic range between methods***

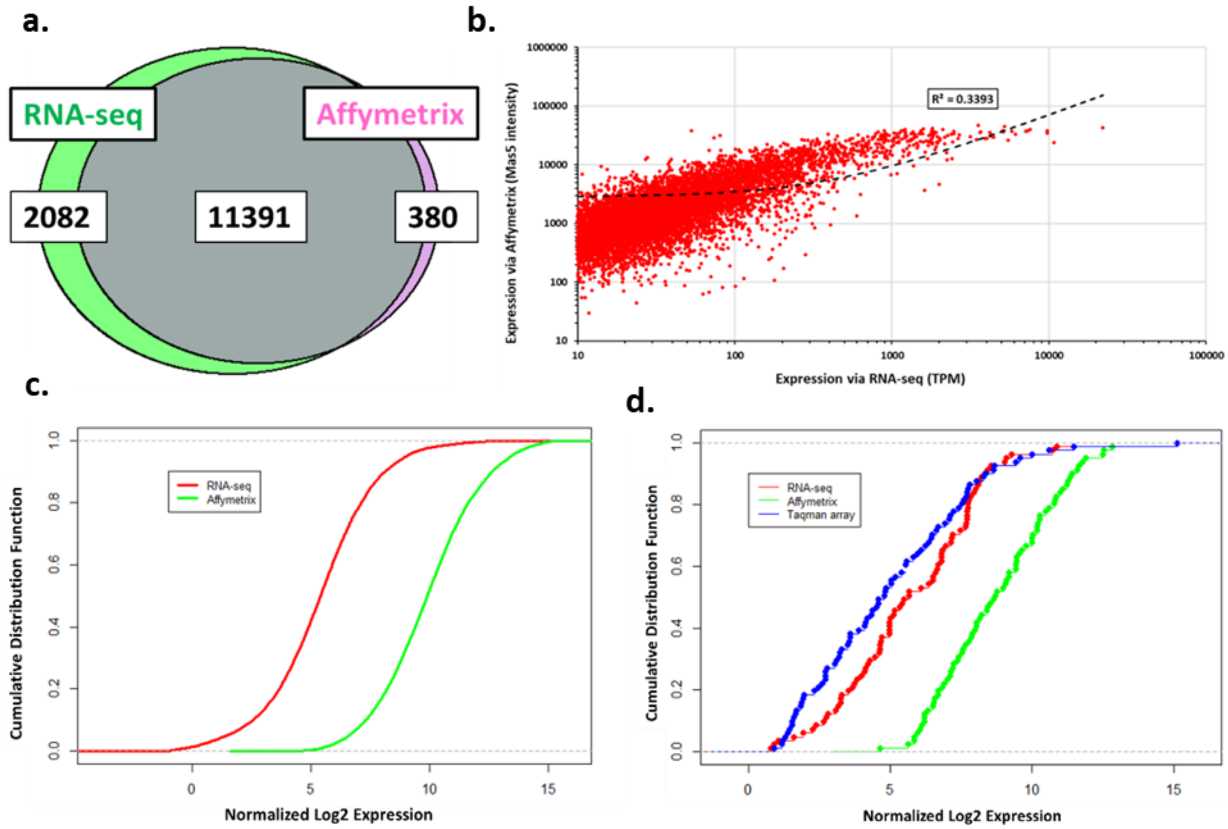


**Figure S3. Gene expression and dynamic range of detection by different methods.** **a)** Venn diagram of the detection of all protein coding genes by Affymetrix HG U133plus2.0 arrays and RNA-seq in pancreatic CAFs. **b)** Correlation of expression values for commonly detected genes by both methods. **c)** and **d)** Cumulative distribution functions (CFDs) showing **c)** the dynamic range of RNA-seq and Affymetrix arrays (HG U133 plus2.0) for all genes; **d)** the same as c), but for GPCRs, detected by RNA-seq, Affymetrix arrays and TaqMan arrays.

To determine how the detection of GPCRs compares with that of the expression of all protein coding genes, we compared the dynamic range of detection for those genes by Affymetrix HG U133plus2.0 arrays and RNA-seq. For estimating the dynamic range of detection for GPCRs, we also included TaqMan GPCR arrays. We found (**Figure S3a)** that RNA-seq detects more genes but the two methods largely detect expression of the same genes. The latter finding contrasts with the results for GPCRs and likely results from the relatively low expression of some GPCRs and limited ability of Affymetrix arrays to detect low abundance transcripts. For most other genes/mRNAs, especially those with intermediate or high expression, the two methods perform similarly, although the magnitude of expression of commonly detected genes does not correlate well between the two methods **(Figure S3b)**.

**Figure S3c** shows the dynamic range of detection for all transcripts by RNA-seq and Affymetrix arrays; gene expression was normalized to log2 units (i.e., log2 of TPM expression for RNA-seq and MAS5 intensity for Affymetrix arrays). We added a small constant to RNA-seq expression values, so that the log2 of the highest expression value in TPM = log2 of the highest expression value in MAS5 intensity. To evaluate the dynamic range for detection by each method, we computed the cumulative distribution function (CDF) for each set of expression values. RNA-seq had a larger detection range (~50-fold) and could detect genes ranging from ~15 to ~ 0 adjusted log2 TPM whereas Mas5 log2 intensities ranged from ~15 to ~5. **Figure S3d** shows a similar trend for GPCRs with respect to the dynamic range for RNA-seq, TaqMan arrays and Affymetrix arrays. GPCR expression was normalized as log2 units (as above) with a small normalization factor added to TaqMan array data and RNA-seq data, so that log2 expression of GAPDH (in all three datasets as a housekeeping gene) was equal and served as the first point (from the right) on the CDF. As the highest expressed gene that was commonly detectable across all three platforms, GAPDH was used as the housekeeping gene for this comparison. The range of expression levels between GAPDH and the lowest-detectable level of GPCR expression allows us to define and compare the dynamic range for each assay. TaqMan arrays and RNA-seq had a similar dynamic range, consistent with the high degree of correlation between the two methods; the dynamic range was lower with Affymetrix arrays.

**Supplemental Tables**

**Table S1.** Types and sources of data used

| **Sample Type** | **Platform** | **Data source** |
|---|---|---|
| Pancreatic Cancer-associated Fibroblasts (CAFs) | • RNA-seq, TaqMan GPCR arrays, Affymetric arrays<br>• Additional samples with RNA-seq and TaqMan GPCR arrays | Insel lab |
| ASPC-1 pancreatic cancer cell line | • TaqMan arrays (Insel lab)<br>• Affymetrix arrays and RNA-seq | Insel lab<br>CCLE |
| Various other cancer cell lines | • Affymetrix arrays and RNA-seq | CCLE |
| Pulmonary artery smooth muscle cells | • RNA-seq and TaqMan GPCR arrays | Insel lab |
| Ventricular cardiac fibroblasts | • RNA-seq and TaqMan GPCR arrays | Insel lab |
| Ovarian cancer (OV) and Lung Squamous Cell Carcinoma (LUSC) tumor tissue | • Affymetrix arrays and RNA-seq | TCGA |

**Table S2. GPCR detection by TaqMan GPCR arrays, Affymetrix HgU133 plus2.0 and RNA-seq**

**Top**: The maximum possible number of GPCRs detectable by each assay. **Bottom**: the number of GPCRs detected by each method in pancreatic CAFs; data shown for 1 CAF replicate (CAF3) as an example.

| **Assay** | **GPCRs with endogenous ligands (endo-GPCRs)** | **Taste receptors** | **Olfactory receptors** | **Opsins** |
|---|---|---|---|---|
| TaqMan GPCR array | 340 | 1 | 7 | 3 |
| Affymetrix Hg U133 plus2.0 | 354 | 20 | 109 | 5 |
| RNA-seq (Allows abundance estimation of any/all annotated genes) | 391 (~400, including several suspected pseudogenes) | 28 | ~850 (includes many suspected pseudogenes) | 10 |

| **Assay** | **GPCRs with endogenous ligands (endo-GPCRs)** | **Taste receptors** | **Olfactory receptors** | **Opsins** |
|---|---|---|---|---|
| TaqMan GPCR array | 111 | 0 | 2 | 1 |
| Affymetrix Hg U133 plus2.0 | 95 | 2 | 10 | 1 |
| RNA-seq (Allows abundance estimation of any/all annotated genes) | 132 | 7 | 20 | 2 |

**Table S3.** qPCR primer sequences

| | |
|---|---|
| *BDKRB1-F* | AATGCTACGGCCTGTGACAAT |
| *BDKRB1-R* | ATTTCTGCCACGTTCAGTTGC |
| *EDNRB-F* | GTCCCAATATCTTGATCGCCAG |
| *EDNRB-R* | AAGGCACCAGCTTACACATCT |
| *CHRM2-F* | ACACCCTCTACACTGTGATTGG |
| *CHRM2-R* | GTCCGCTTGACTGGGTAGG |
| *AODRA1-F* | CCACAGACCTACTTCCACACC |
| *AODRA1-R* | TACCGGAGAGGGATCTTGACC |
| *BDKRB2-F* | CCGAAAGAAGTCTTGGGAGGT |
| *BDKRB2-R* | CTGGCGTTCCACGGAGATG |
| *GPR115-F* | TTTAAGGACTCAACTGGTGCATC |
| *GPR115-R* | ACACTCTCAATGGTCTCTGGAG |
| *GPRC5B-F* | CACGCCCAACTACTTCGACA |
| *GPRC5B-R* | AGCTGCATTGTGTTCATCCAT |
| *EMR2-F* | AGAAGCAAGTAGACAGGAGTGT |
| *EMR2-R* | TTCTGTGCCTGATTCCAGTCG |
| *ADRA2A-F* | TCGTCATCATCGCCGTGTTC |
| *ADRA2A-R* | AAGCCTTGCCGAAGTACCAG |
| *CELSR1-F* | GGCGTTGTTTGAGAACGAACC |
| *CELSR1-R* | AGAGTCGATTCGGAAGTAGCC |
